# Defining the Substrate Envelope of SARS-CoV-2 Main Protease to Predict and Avoid Drug Resistance

**DOI:** 10.1101/2022.01.25.477757

**Authors:** Ala M. Shaqra, Sarah Zvornicanin, Qiu Yu Huang, Gordon J. Lockbaum, Mark Knapp, Laura Tandeske, David T. Barkan, Julia Flynn, Daniel N.A. Bolon, Stephanie Moquin, Dustin Dovala, Nese Kurt Yilmaz, Celia A. Schiffer

## Abstract

Coronaviruses, as exemplified by SARS-CoV-2, can evolve and spread rapidly to cause severe disease morbidity and mortality. Direct acting antivirals (DAAs) are highly effective in decreasing disease burden especially when they target essential viral enzymes, such as proteases and polymerases, as demonstrated in HIV-1 and HCV and most recently SARS-CoV-2. Optimization of these DAAs through iterative structure-based drug design has been shown to be critical. Particularly, the evolutionarily conserved molecular mechanisms underlying viral replication can be leveraged to develop robust antivirals against rapidly evolving viral targets. The main protease (M^pro^) of SARS-CoV-2, which is evolutionarily constrained to recognize and cleave 11 specific sites to promote viral maturation, exemplifies one such target. In this study we define the substrate envelope of M^pro^ by determining the molecular basis of substrate recognition, through nine high-resolution cocrystal structures of SARS-CoV-2 M^pro^ with the viral cleavage sites. These structures enable identification of evolutionarily vulnerable sites beyond the substrate envelope that may be susceptible to drug resistance and compromise binding of the newly developed M^pro^ inhibitors.

## INTRODUCTION

Coronaviruses can jump from animal reservoirs to humans and cause major outbreaks, which has happened at least three times (1–3) during the past two decades. Such zoonotic transmissions are more common than realized (4, 5), where the virus evolves to acquire the ability to infect humans and then further adapt to human host. This ability to mutate presents a challenge to the longevity of vaccine efficacy as well as any potential antiviral therapeutics. To combat current and future coronavirus pandemics, a combination of preventive and therapeutic options is needed, including direct-acting antivirals (DAAs) to treat vulnerable populations to decrease morbidity and mortality.

Coronaviruses are non-segmented positive-sense single-stranded RNA viruses. The viral genome encodes open reading frames two of which are translated into polyproteins that must be cleaved to release individual proteins. Processing the majority of these sites, including autocleavage, is the function of the coronavirus main protease, M^pro^. Because cleavage of these sites to liberate the viral proteins is essential for replication of the virus, any intervention that stops this process, whether be a mutation or an inhibitor, would block viral growth. M^pro^ has also been implicated in cleaving sites in key cellular host factors to likely enhance viral replication (6). Due to the essential function of SARS-CoV-2 M^pro^, this protease has been targeted with small molecule inhibitors with currently one molecule FDA-authorized as a clinical drug and several others in the developmental pipeline (7). Targeting viral proteases with DAAs has demonstrated great success in HIV-1 and HCV infections (8–11), and initial results on inhibitors of M^pro^ are extremely promising to become critical therapeutics against SARS-CoV-2.

The substrate cleavage sites processed by SARS-CoV-2 M^pro^ have a conserved glutamine after which the protease cleaves but are otherwise diverse in amino acid sequence, especially at the prime side (C-terminal to the cleavage). As we have found for HIV-1 and HCV NS3/4A proteases previously (12–14), the viral proteases bind substrates of diverse amino acid sequences through a conserved 3D structure or shape for which we coined the term *substrate envelope*. In addition to the molecular basis of specificity and recognition, the substrate envelope effectively explains susceptibility of protease inhibitors to resistance (15, 16). Active site residues that contact the inhibitor outside the substrate envelope can mutate without affecting substrate recognition to confer resistance. In contrast, inhibitors that fit well within the substrate envelope optimally occupy the same space as the natural substrates thus leveraging the evolutionary constraint of substrate recognition.

To define the structural basis of substrate recognition and thereby determine the envelope to avoid drug resistance, we have determined 9 substrate-cocrystal structures of SARS-CoV-2 M^pro^. The high-resolution structures reveal the intermolecular interactions essential for molecular recognition and enable defining the conserved substrate envelope. An additional 6 product cocrystal structures bound to the cleaved N-terminal side of the substrate peptide were also determined and comparatively analyzed. The substrate envelope maps out the specificity of M^pro^ and reveals intermolecular interactions that are essential for enzymatic function. These interactions will guide the design of inhibitors and help pinpoint which M^pro^ residues are vulnerable to the occurrence of resistance.

## RESULTS

The crystal structures of SARS-CoV-2 M^pro^ with 9 substrate and 6 product complexes were determined to sub-2.0 Å resolution (**Table S1 & Table S2**). There is little sequence conservation among the natural cleavage sequences (**Figure 1A**), except the fully conserved glutamine at P1 and a hydrophobic L/F/V at the P2 position. The 12-mer peptides, corresponding to the cleavage sites, were largely ordered from P5-P2′ in the cocrystal structures. As in previous apo and inhibitor-bound crystal structures (17–20), M^pro^ crystallized as a homodimer (**Figure 1B**). Six of the nine complexes were solved with the dimer in the asymmetric unit (in P2_1_ or P2_1_2_1_2_1_), with both active sites in the homodimers occupied with the substrate, while the other three had a monomer in the asymmetric unit (in C2_1_). As was previously observed, the N terminal serine residue of one monomer was reaching into the active site of the other monomer, completing the S1 pocket of the other monomer’s active site (**Figure 1C**).

**Figure 1.**
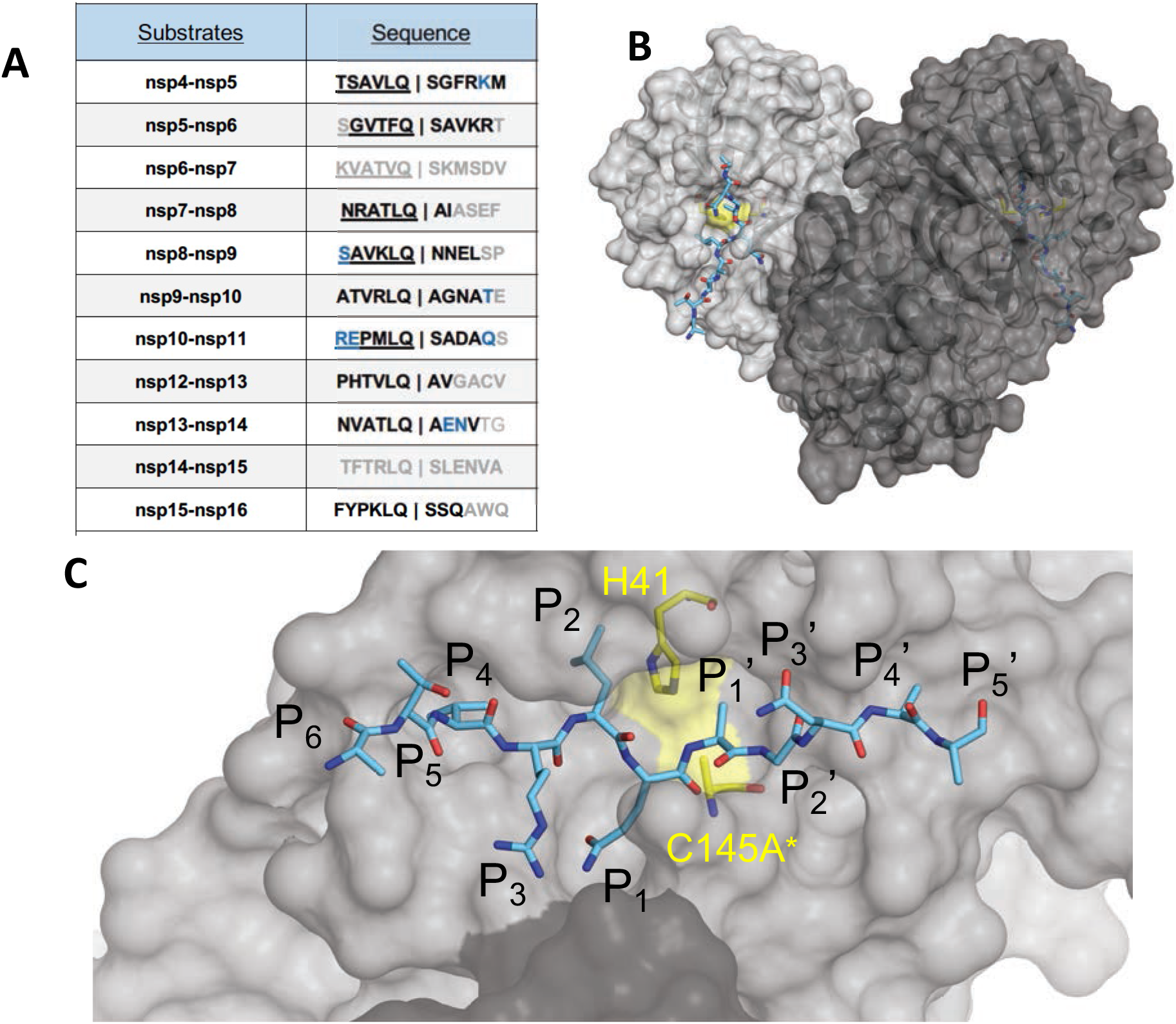
The amino acid sequences and binding of substrates to SARS-CoV-2 M^pro^ active site. (**A**) Viral polyprotein cleavage sites processed by M^pro^ to release non-structural proteins (nsp). The one-letter amino acid codes of cleavage site sequences, where bold letters indicate fully resolved residues and blue are stubbed side chains in the cocrystal structures. Underlined N-terminal sequences correspond to product complexes with independently determined cocrystal structures. (**B**) Crystal structure of SARS-CoV-2 M^pro^ with a substrate peptide (nsp9-nsp10) bound at the active site of both monomers (light and darker gray). The peptide is depicted as cyan sticks and the catalytic dyad is colored yellow. (**C**) Close-up view of one of the active sites in panel B, with the protease in surface representation. The asterisk indicates catalytic cysteine was mutated to prevent substrate cleavage. The cleavage occurs between positions P1 and P1′.

In these cocrystal structures, both active sites were fully occupied with the noncleaved substrate in essentially the same conformation. This is in contrast to the “half-site” activity previously suggested for SARS-CoV-1 M^pro^ (21) which was adopted to SARS-CoV-2 partly based on a proposed inactive conformation of Glu166 in one of the monomers in an inhibitor-bound crystal structure (19). In all our structures, Glu166 is in the same conformation in both active sites. Thus, our structures support that both monomers can be simultaneously active.

In all the current cocrystal structures (**Figures S1 and S2**), the substrate peptide was extended along the M^pro^ active site with the scissile bond between the P1 glutamine and P1′ residue positioned between the catalytic dyad. Although M^pro^ cleaves these sites in the context of a longer polypeptide to release viral non-structural proteins (nsps), molecular interactions with the protease are localized to residues proximal to the scissile bond. The N-terminal (or non-prime) side of the substrate had an antiparallel beta-strand conformation which was conserved in all structures, with the substrate residues and side chains well resolved. The binding mode of the C-terminal residues (prime side) was more varied among the substrates and certain residues, especially beyond P3′ position, and lacked full electron density. In addition to the fully conserved P1 glutamine, which is stabilized through multiple molecular interactions, the large hydrophobic residue at P2 was extended deep into the S2 binding pocket in all structures.

### Hydrogen bonding network that ensures M^pro^ substrate specificity

To investigate how substrate binding is stabilized at the M^pro^ binding site and identify conserved features, we analyzed the intermolecular interactions between the substrates and M^pro^ active site residues. The substrate peptides and the active site residues formed multiple networks of hydrogen bonds that stabilized the binding interaction (**Figure 2A**) including some mediated by conserved waters. In all of the cocrystal structures, the catalytic His41 was stabilized by one such network where the conserved, potentially catalytic water is coordinated by Asp187 and His164 (**Figure 2B**). The sidechain of the conserved P1 glutamine was also extensively coordinated in another hydrogen bond network. The first shell of this network includes the sidechains of His163 and Glu166, the backbone of Phe140, and three conserved waters. This network was further stabilized by the sidechain of Asn142 which coordinates the conserved waters and Ser1 from the other monomer, stabilizing the position of both the backbone of Phe140 and the sidechain of Glu166. This extensive network underlies the requirement of homodimer formation in defining the P1 specificity for glutamine.

**Figure 2.**
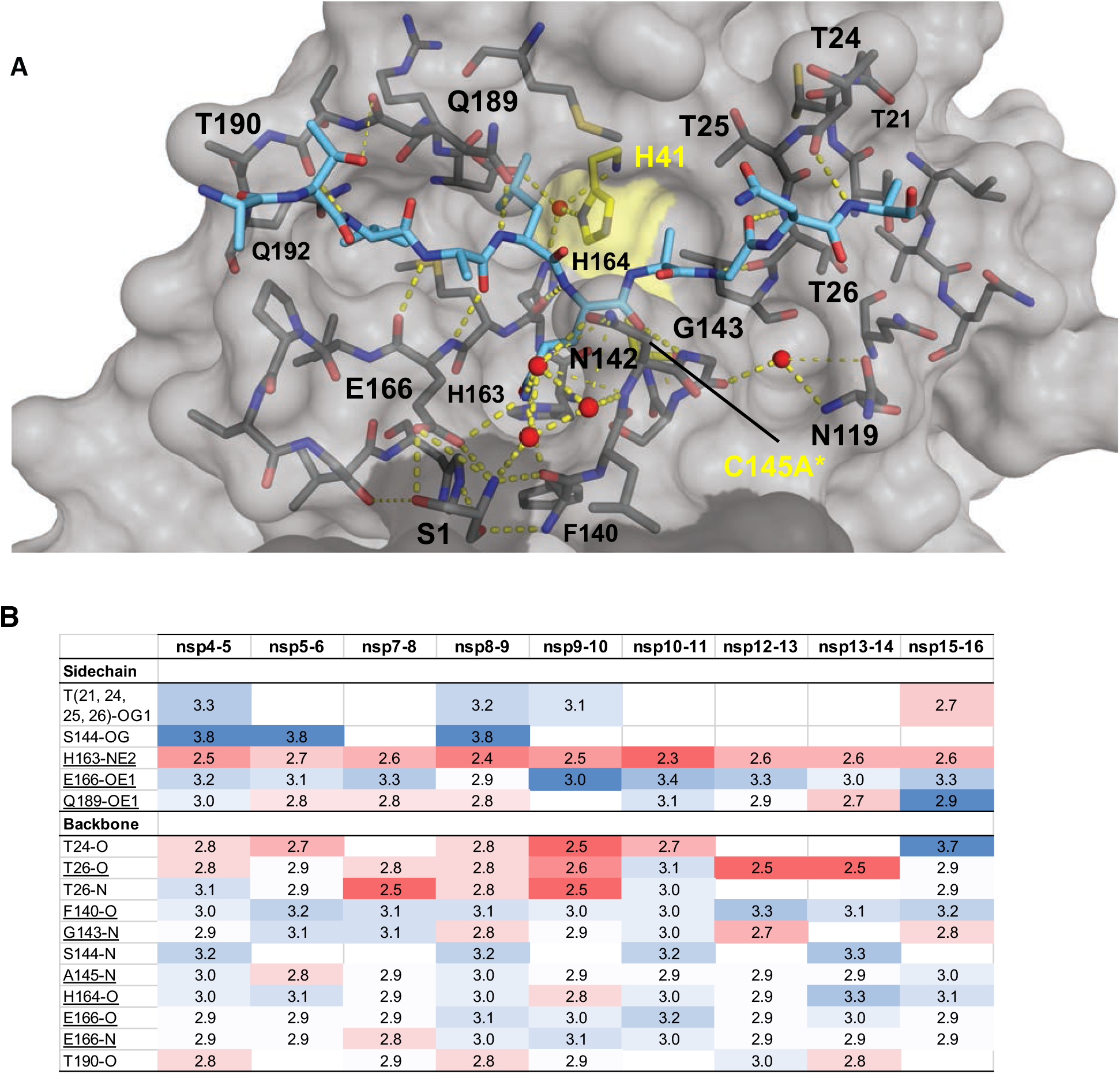
Intermolecular hydrogen bonds in M^pro^ substrate cocrystal structures. **(A)** Hydrogen bonds between bound nsp9-nsp10 substrate and M^pro^. The substrate peptide is depicted as cyan sticks and the protease is in gray surface representation with the catalytic dyad colored yellow. Yellow dashed lines indicate hydrogen bonds (thicker lines for stronger bonds with distance less than 3.5 Å) and red spheres denote conserved water molecules. Serl depicted as sticks belongs to the other monomer (shown in darker gray). **(B)** Hydrogen bonds that are conserved in three or more substrate complexes, underlined completely conserved, top interacting with M^pro^ sidechains and bottom with M^pro^ backbone atoms, color coded by the closeness of the hydrogen bond.

Beyond the P1 sidechain, the conserved hydrogen bonds were with the backbone of the substrates, largely in the form of backbone-backbone hydrogen bonds. The N-terminal side of the P1O was coordinated by three nitrogen atoms, Gly143N, Ser144N and Cys(Ala)*145N, while P1N:His164O, P2O:Gln166N, P3N:Gln166O and P4N:T190O stabilized the peptide. Only the sidechain of Gln189 also hydrogen bonded to the P2N. On the C-terminal (primed) side of the substrates, the interactions were again backbone-backbone hydrogen bonds including P1′O:Gly143N; P2′N:Thr26O; P2′O:Thr26N; P4′N:Thr24O. Each cleavage site was further stabilized by additional sequence-specific hydrogen bonds, often coordinating highly ordered water molecules.

### Packing of diverse substrates is largely conserved with M^Pro^

To analyze packing of active site residues around the substrate peptides and quantify the inter-molecular interactions, van der Waals (vdW) contacts were calculated for each protease-substrate pair (**Figure 3A**). Overall, the contact pattern was consistent among the substrates, where M^Pro^ residues at the S3-S2′ sites contributed significant vdW contacts with the substrate (**Figure 3B**). At the S1 subsite, the conserved P1 glutamine had substantial vdW interactions with Asn142, consistent with the extensive hydrogen bonding network described above regarding P1 specificity. Significantly, the sidechain of Gln189 forms a cavity which engulfs the P2 residue and forms the most extensive vdW contact for each substrate. His164, Met165, and Glu166 also form a pocket that accommodates the P4 residue with extensive contacts. As prime side residues are poorly conserved between M^Pro^ substrates, the enzyme residues that contribute significant vdW contacts differ. However, several vdW contacts were also conserved on the prime side; the threonine cluster of Thr24, Thr25, and Thr26 hydrogen bonds to stabilize prime side residues prior to substrate cleavage. While not the most extensive, the vdW contacts of catalytic dyad His41 and Cys145A* were highly conserved and consistent between all nine substrates. Analyzing the packing of the substrates, the interactions largely mirror the packing of the enzyme with the conserved Gln P1 packing the most extensively, followed by P2 and P4, and then P1’ (**Figure S3**). Each of these sites make some contact with conserved residues (**Figure 3C**). Overall, despite the vast variation in substrate amino acid sequences, the packing around the bound substrates and interactions of protease residues were highly structurally conserved.

**Figure 3.**
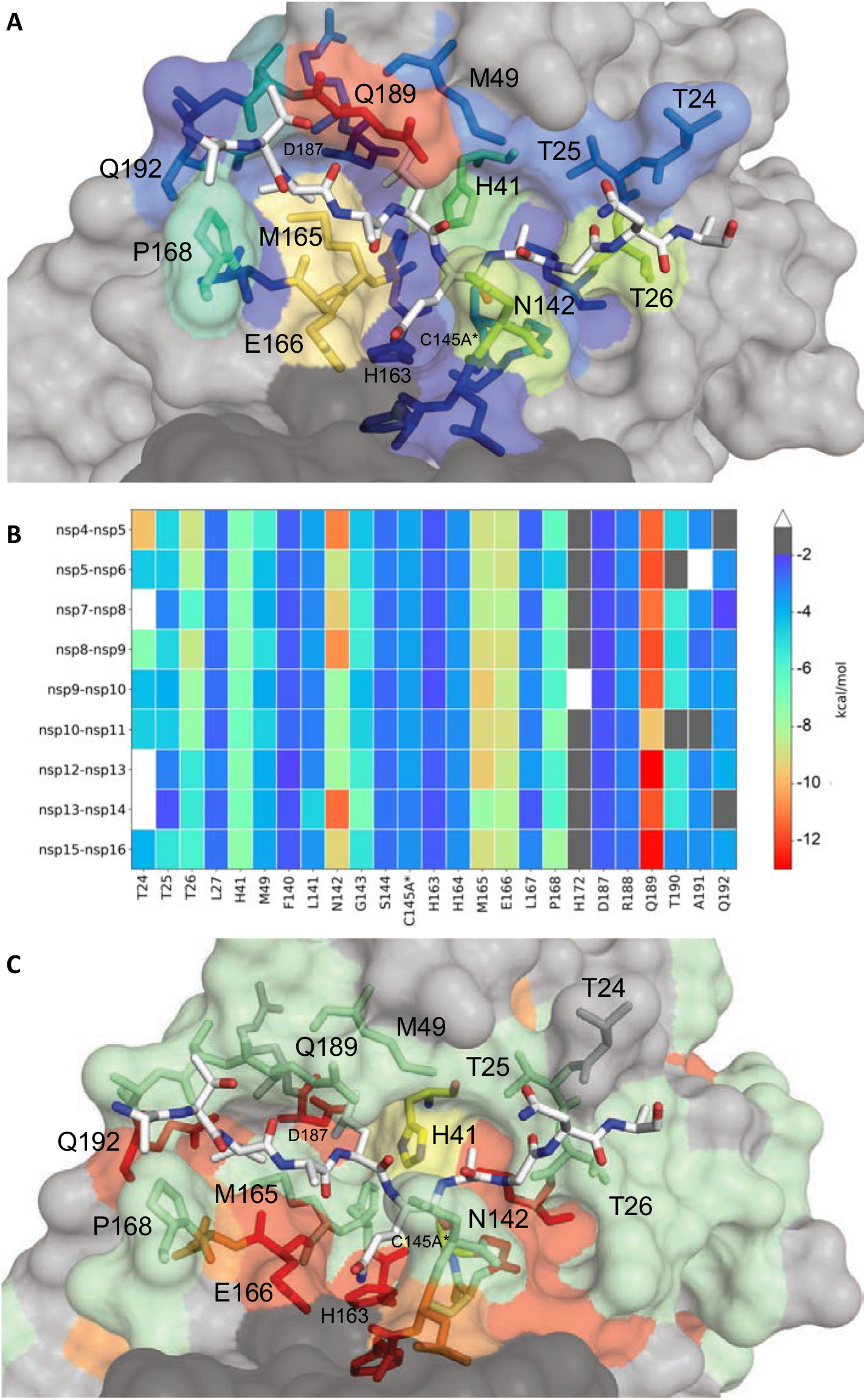
Extent of substrate interactions and conservation of M^pro^ surface residues. (**A**) Close-up view of the nsp9-nsp10 substrate bound to M^pro^ active site in the cocrystal structure where the substrate peptide is depicted as white sticks and the protease is in surface representation. The protease residues are colored according to the extent of van der Waals interactions with the substrate, with warmer colors indicating more interaction. (**B**) Conservation of substrate-protease van der Waals interactions among the 9 cocrystal structures determined. Heat map coloring by extent of van der Waals contact by residue. **(C)** Amino acid sequence conservation of M^pro^ between 7 (**Figure S4**) coronaviral species depicted on the structure where surface residues conserved in all 7 (red), 5-6 (orange), 3-4 (green) and less than 3 (highly variable; gray) sequences are indicated by color.

Interestingly among coronaviruses little sequence conservation exists within M^pro^, even around the active site, indicating that most of the residues that line the surface of the active site groove tolerate quite a bit of variability (**Figures 3C and S4**), consistent with saturating mutational analysis as performed by Flynn et al. (22). These include residues Asn142, Met165, Glu166 and Gln189 which form extensive vdW interactions and hydrogen bonds with the substrates (**Figures 2 and 3**). In the mutational analysis of Flynn et al (22) in the SARS-CoV-2 background, Asn142, Glu166 and Gln189 all tolerate extensive variability with Met165 being less tolerant; between coronaviral species Asn142 (Cys and Ala), Met165 (Leu and Ile), and Gln189 (Glu and Pro) are variable while Glu 166 is conserved. In the M^pro^ structure, Gln189 is located in the center of the structurally and sequence-wise variable loop that is anchored by two completely invariant residues Asp187 and Gln192. Another major exception to the variability is His163 which appears to be intolerant to change and coordinates the P1 glutamine. Taken together the surface of M^pro^ can tolerate extensive variation while maintaining activity and structural interactions.

### The M^pro^ substrate envelope

The broad range of sequences M^pro^ cleaves (**Figure 1**) present a challenge of molecular recognition for this viral protease in facilitating viral maturation. To elucidate how this is achieved, the substrate complex structures were superimposed based on a set of invariant active site residues within one monomer (see Methods). The substrate structures superimpose very well, especially the P2-P1′ residues (**Figure 4**). Structurally, the M^pro^ enzyme in complex with nsp5-nsp6, to accommodate the unique Phe at P2, diverged the most from the rest of the complexes. The divergence was especially in the loop that closes over the substrates from Asp187 to Gln192, where the Gln189 was shifted by 1.0 Å relative to the other complexes. This adaptability of this loop appears to be key to accommodate the diverse sequences.

**Figure 4.**
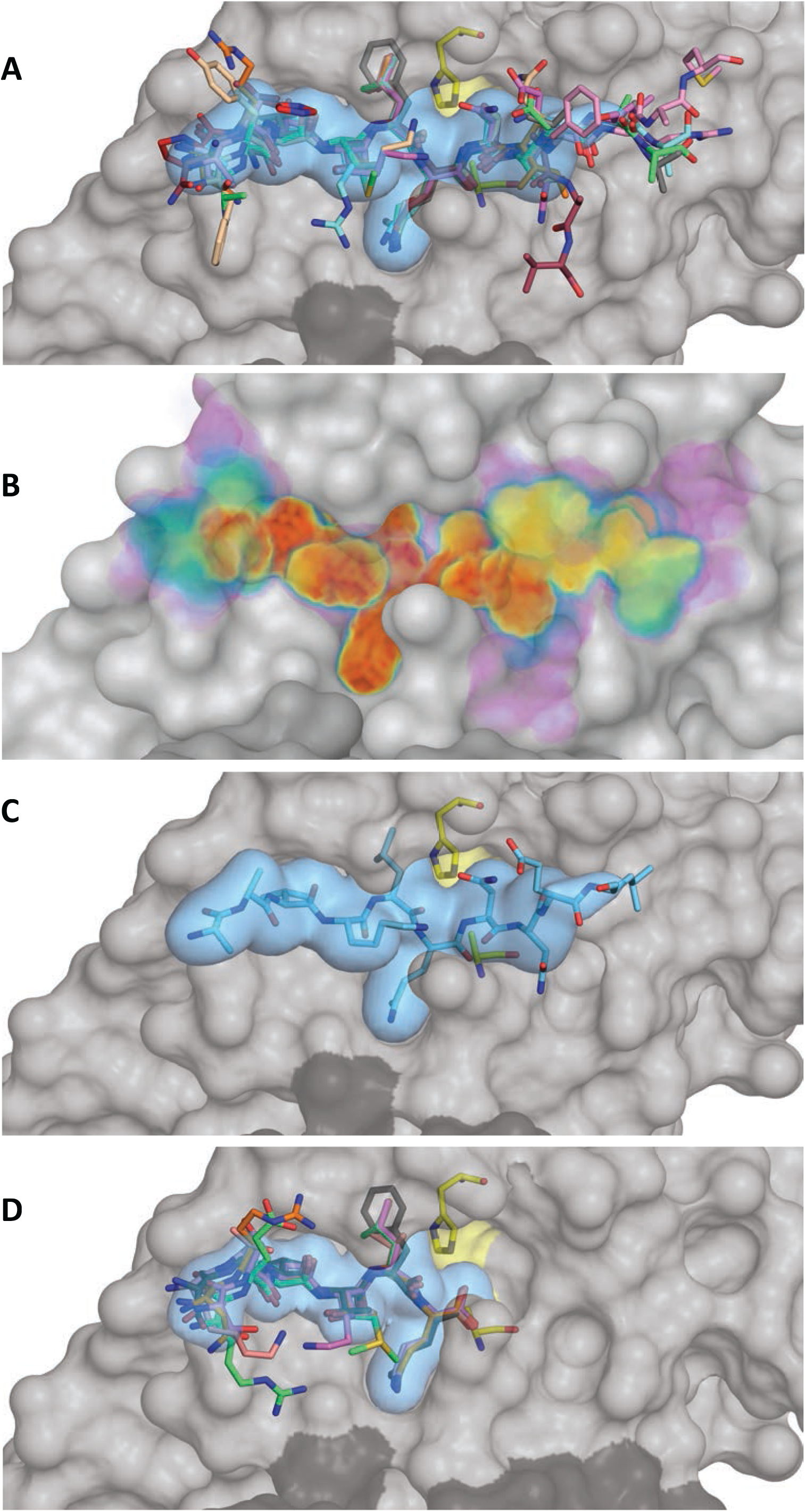
The substrate envelope of SARS-CoV-2 M^pro^. (**A**) The substrates bound at the M^pro^ active site are depicted as sticks in the superimposed cocrystal structures, where the protease is in gray surface representation. The consensus volume occupied by the substrates define the substrate envelope, shown as the blue volume and is the intersection of any four of the nine substrates. (**B**) The gradient substrate envelope colored according to the number of substrates that occupy the consensus volume. Purple to red gradient indicates less to more consensus. (**C**) Substrate nsp8-nsp9 in the substrate envelope. (**D**) Superposition of cocrystal structures with cleaved N-terminal product complexes, defining the product envelope (blue volume).

On the molecular level, what accounts for substrate specificity with such diverse sequences? As we have previously seen with HIV-1 and HCV NS3/4A proteases where substrate shape accounts for specificity, despite differences in sequence (12, 23), M^pro^ appears also to recognize a conserved substrate shape. This shape defines the substrate envelope. The substrate envelope was calculated by overlapping consensus volumes to visualize the space occupied at the M^pro^ active site (**Figure 4A)**. While certain regions, particularly at the C-terminal side, deviate from the consensus these moieties are largely solvent exposed. To comprehensively evaluate the consensus and conservation of the occupied volume at the M^pro^ active site, we also calculated a gradient substrate envelope reflecting how many substrates overlap at a given position (**Figure 4B**). In this gradient envelope, the space occupied by only a single substrate has the lowest score (shown in purple) while the space that all substrates occupy has the highest score (shown in red). As an example, the nps8-nsp9 substrate, which is the most conserved cleavage site between coronaviral species (24), fits extremely well within the substrate envelope (except the unusual P1′ Asn, where the hydrophilic end of the side chain protrudes beyond the envelope) (**Figure 4C**). The volume from P4 to P2′ is highly conserved between all of the substrate complexes, despite the variation in amino acid sequences; this high conservation reflects the specificity and likely evolutionarily-constrained regions of the enzyme.

In addition to the full peptide substrates corresponding to the viral polyprotein cleavage sites, we determined six product complexes. The proposed reaction mechanism for M^pro^ involves breakage of the scissile bond and formation of an acyl-enzyme complex with a covalent bond between the N-terminal fragment and catalytic cysteine. Our cocrystals captured the N-terminal product after the cleavage reaction was complete where no covalent bond exists with the catalytic cysteine. All 6 cleaved substrates bound at the N-terminal side of the active site superimposed very well, defining a “product envelope” (**Figure 4D**). There were no major rearrangements or shifts in the backbone and except minor side chain conformers, the products bound similarly to the noncleaved substrates. The nsp5-nsp6 cleavage site with the P2 Phe was once again the outlier where the Phe was in an alternate rotamer and the 187-192 loop was shifted relative to the other complexes (**Figure S5**). Overall, however, the product envelope recapitulated the consensus volume revealed by the substrate envelope for the N-terminal part of the M^pro^ active site.

### Inhibitor fit within the substrate envelope and resistance mutations

Where inhibitors protrude from the substrate envelope is an excellent indicator of susceptibility to resistance mutations, as we and others have shown extensively for multiple targets and various inhibitors (10, 15, 16, 25–28). Thus, our newly determined substrate envelope of M^pro^ provides a predictive tool for both current and potential inhibitors that may become clinical drugs. Sites where inhibitors protrude from the consensus volume to contact protease residues pose vulnerabilities as changes at those locations could weaken inhibitor binding without compromising substrate binding and cleavage.

The binding mode and fit within the substrate envelope of four SARS-CoV-2 M^pro^ inhibitors that are either under emergency authorization by the FDA or currently in advancing development. These inhibitors include two covalent inhibitors developed by Pfizer, PF-07321332 (29) (emergency use) (**Figure 5A**) and PF-00835231 (30) (**Figure 5B**), a non-covalent inhibitor compound 21 (31) (**Figure 5C**), and the most potent covalent inhibitor from the COVID Moonshot project, compound 11 (32) (**Figure 5D**). All the covalent inhibitors span P4-P1 portion of the actives site, while the non-covalent inhibitor, compound 21(31), extends into the P1’ region. The variation in contact of the inhibitors with the active site can be seen in the variation in vdW contact where Met 165 and Glu 166 make the most extensive interactions (**Figure S6**). These four inhibitors all have similar vulnerabilities as assessed by the substrate envelope; the lactam ring and other ringed moieties protrude out at P1 near Asn142 and Glu166 while at P2 inhibitors protrude along the conformationally variable loop from 187-192 and Gln189. Three residues appear very flexible adopting varied conformations depending on the inhibitor: Asn142, Gln189 and M49; in contrast with their invariant conformations when bound to substrates. These variations in conformation often occur near where the inhibitors protrude from the M^pro^ substrate envelope. These sites are not conserved among coronaviral species and tolerate mutations (22), rendering these residues potential sites for resistance. Additionally, although the conformation of Glu166 is conserved structurally, this residue also appears to tolerate mutations (22) thus potentially can also confer resistance if variation occurs. As often occurs when resistance arises, at all 4 sites (residues 49, 142, 166 and 189) a change in inhibitor interactions, either by sidechain becoming more rigid (His, Tyr, Trp or Phe) to cause a clash; becoming smaller (Ala, Thr, Val, Ser) to cause a loss of contact or a change in charge Glu ➔ Gln, could dramatically alter drug binding, while maintaining the substrate envelope and recognition.

**Figure 5.**
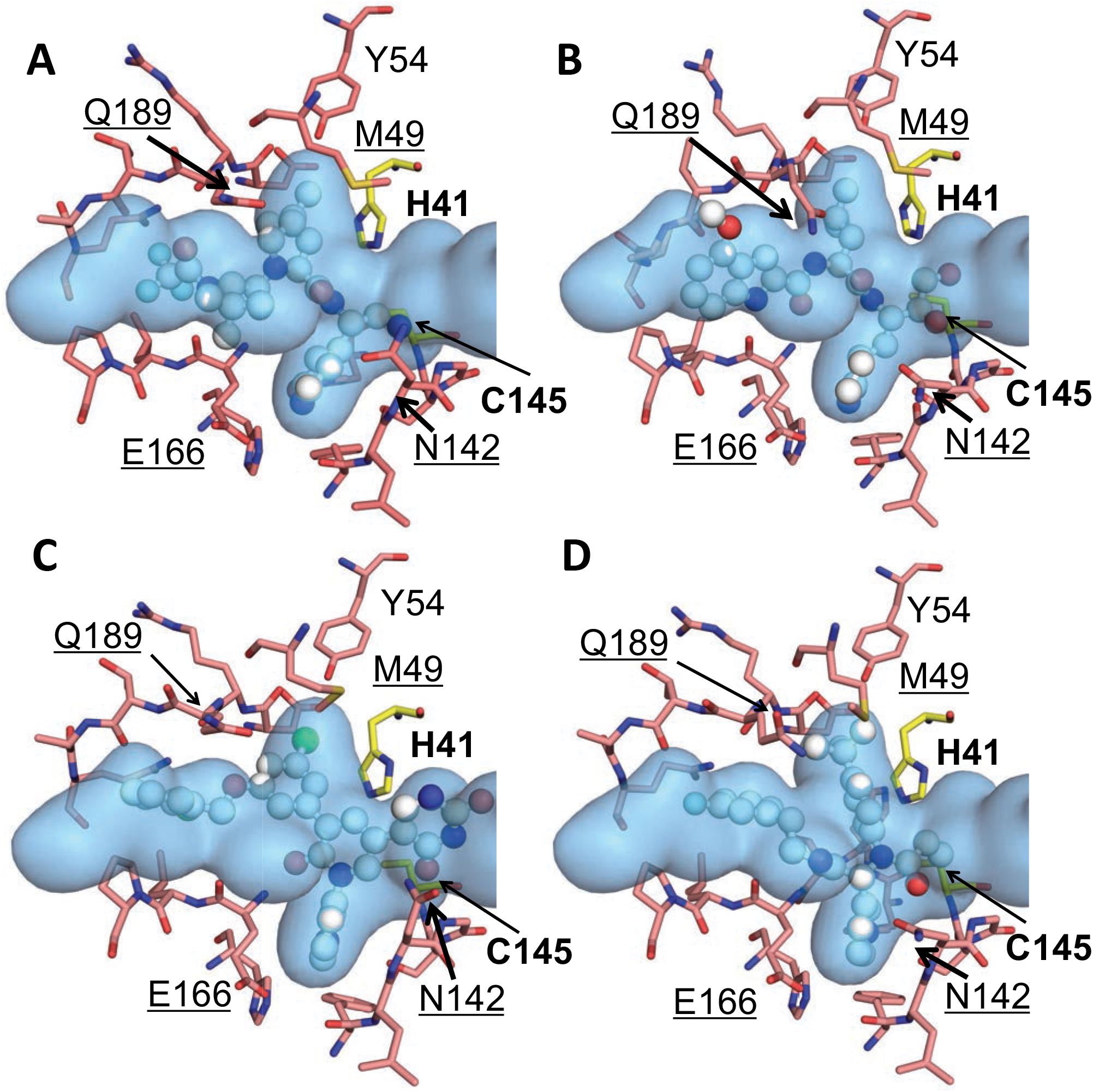
The fit of protease inhibitors within the substrate envelope. **(A)** PF-07321332 (25), PDBID:7RFS **(B)** PF-00835231 (26), PDBID: 6XHM **(C)** Noncovalent potent compound 21 (27), PDBID: 7L13 **(D)** Moonshot compound 11 (28), PDBID: 7NW2. The inhibitors are in ball-and-stick representation and the substrate envelope is depicted as the blue volume. The catalytic dyad residues are labeled in bold while the underlined labels are for the contact residues that interact with one or more inhibitor and may be vulnerable to resistance.

## DISCUSSION

COVID-19 pandemic caused by SARS-CoV-2 resulted in the fastest developed DAAs getting to patients ever. While this is a great accomplishment, the quick evolving of SARS-CoV-2 with variants of concern and the likelihood of other pandemic potential coronaviruses arising present a challenge to developing an arsenal of drugs. Preemptive strategies need to be developed to design inhibitors that are less susceptible to drug resistance and escape to ensure their longevity and efficacy both in the current outbreak and in the future for pan-CoV DAAs.

One critical drug target has been the SARS-CoV-2 viral protease Mpro where there has been a substantial effort to design DAAs to thwart the virus. Initial efforts by both academia and industry have successfully identified quite a variety of inhibitors through a combination of structure based and medicinal chemical approaches (19, 33–36) The vast majority of these Mpro inhibitors are covalent targeting the catalytic cysteine, but several promising noncovalent inhibitors with low nanomolar potency are in development. The first Mpro inhibitor that has made it through clinical trials and received emergency authorization is PF-07321332 (29). Now known as nirmatrelvir and formulated together with ritonavir to prevent degradation, the clinical drug Paxlovid decreases the chance of hospitalization in patients infected with SAR-CoV-2 by 95%. While this is incredibly promising, the possibility that SARS-CoV-2 evolves in such a way that this efficacy is compromised, or another CoV arises that makes PF-07321332 less potent urges the design of novel Mpro inhibitors that preemptively avoid resistance.

In this study we have defined the substrate envelope of SARS-CoV-2 Mpro, which reveals the molecular basis of substrate specificity and can guide the design of active site inhibitors to avoid drug resistance. By determining 15 cocrystal structures, with 9 corresponding to the diverse viral substrate cleavage sites and 6 with N-terminal products, we were able to identify the conserved features required to recognize the diverse substrate sequences and define the substrate envelope. Specific interactions with protease active site residues as well as a conserved network of water molecules ensure substrate specificity and proper geometry. The C-terminal side of the substrates were more divergent in their binding modes and had weaker inter-molecular interactions, consistent with the reaction mechanism and the N-terminal side being captured in the product cocrystal structures. In addition to molecular recognition of substrates, the cocrystal structures enabled defining the viral substrate envelope. The conserved substrate envelope defines the molecular interactions that underlie the requirement for proper processing of the viral polypeptides, and thereby imposes an evolutionary constraint on the survival of the virus. As such, the virus cannot evolve mutations that disrupt these conserved interactions without compromising viral survival.

As we have previously demonstrated, where inhibitors protrude outside the substrate envelope to establish interactions with mutable residues correlate with the sites of resistance mutations. As we have shown in **Figure 5** many of the most promising inhibitors have interactions beyond the substrate envelope at residues Met49, Asn142, Glu166 and Gln189. Variation at these sites both within coronaviral variants (**Figure 3C and SM**) and comprehensive saturation mutagenesis analysis in Flynn et al (22) reveal that these active site residues are tolerant to mutations; thus leaving the inhibitors, including nirmatrelvir, vulnerable to viable mutations which would confer drug resistance.

As the COVID-19 pandemic continues into a third year, second and third generation SARS-CoV-2 DAAs are necessary both to thwart the rapid evolution of variants of concern and to prepare for future coronaviral pandemics. Development of additional M^pro^ inhibitors is essential to target and stop such variants, as the enzyme is evolutionary constrained by the necessity to cleave 11 viral sites. As we have previously demonstrated in HIV and HCV, potent protease inhibitors that fit within the substrate envelope are less prone to resistance as a mutation impacting these inhibitors will simultaneously impact the protease’s ability to recognize viral substrates. While the prevalence of clinical resistance may depend on various factors, constraining inhibitors within the substrate envelope to leverage conserved biological features is a powerful strategy to curb evolution and prolong the longevity of the next generation DAA M^pro^ inhibitors.

## METHODS

### Expression and purification of SARS-CoV-2 M^pro^

His6-SUMO-SARS-CoV-2 M^pro^(C145A) was cloned into a pETite vector. Hi-Control BL21(DE3) *E. coli* cells were then transformed with this vector using standard techniques. A single colony was used to start an overnight culture in LB + kanamycin media. This culture was used to inoculate 2 x 1 L cultures in TB, supplemented with 50 mM sodium phosphate pH 7.0 and 50 μg/mL kanamycin. These cultures grew in Fernbach flasks at 37 °C while shaking at 225 rpm, until the OD600 reached approximately 1.5, at which point the temperature was reduced to 19 °C and 0.5 mM IPTG (final) was added to each culture. The cells were allowed to grow overnight. The cell mass was then resuspended in IMAC_A buffer (50 mM Tris pH 8.0, 400 mM NaCl, 1 mM TCEP) prior to lysis by three passes through a cell homogenizer at 18,000 psi. Cell lysate was then clarified by centrifugation at 45,000 x g for 30 minutes. Clarified lysate was flowed through a 5 mL Ni-Sepharose excel column on an AKTA FPLC. The column was pre-equilibrated with 5 CV of IMAC_A. The material was flowed using a sample pump with a flow rate of 5 mL/min. Following column loading, the column was washed with IMAC_A buffer until the A280 stabilized, at which point it was reset to 0. The material was then slowly eluted with a linear gradient of IMAC_B (50 mM Tris pH 8.0, 400 mM NaCl, 500 mM imidazole, 1 mM TCEP) over 40 column volumes. The presence of M^pro^ in the elution peak was confirmed by ESI-LC/MS. The SUMO tag was then cleaved by addition of ULP1 to the pooled fractions from the IMAC purification, resulting in an authentic N-terminus. Cleavage proceeded at room temperature overnight while dialyzing into 3 L of IMAC_A Buffer via 10,000 MWCO dialysis cassette. The protein was then flowed over 5 mL of Ni-NTA resin pre-equilibrated with IMAC_A buffer to remove the cleaved tag. The remaining protein was “pushed” out of the resin with an additional 5 mL wash with IMAC_A buffer. Protein from the rIMAC purification was concentrated to approximately 3 mL prior to purification via SEC. A Superdex 75 16/60 column was pre-equilibrated with fresh SEC Buffer and the protein was flowed through the column at 1 mL/min while collecting 1.5 mL fractions. Fractions in the included peak were pooled and concentrated, then stored at −80 °C. We noticed that during purification the protein behaved better if kept at room temperature. Exposure for long periods of time to lower temperatures (eg: a cold room) mostly led to precipitation.

### Protein Crystallization

Purified SARS-CoV2-M^pro^, the inactive form, C145A, and lyophilized substrate peptides were purchased from ELIM Biosciences and provided by Novartis Institutes for Biomedical Research. M^pro^-NSP substrate and product cocrystals were produced according to conditions previously described by our group (37, 38) with some modifications. 10 mg/mL of protein was thawed on ice and diluted to 6.7 mg/mL in 20 mM HEPES pH 7.5, 300 mM NaCl buffer. Prior to complex formation, the protein was centrifuged at 13,000 *xg* for 1 minute at 4 °C to remove insoluble particulates that may hinder crystal growth. Substrate and product complexes were formed by incubating M^pro^ with 10-fold molar excess of substrate peptides on ice for 1 hour. Crystals were grown using 24-well, pre-greased, VDX hanging-drop trays (Hampton Research Corporation) at various protein to precipitant ratios (1 μL:2 μL, 2 μL:2 μL, and 3 μL:2 μL) with 10-20 % (w/v) PEG 3350, 0.20-0.30 M NaCl, and 0.1 M Bis-Tris-Methane:HCl pH 5.5. Crystal growth took place at room temperature and required 1-2 weeks to obtain diffraction quality cocrystals. In some cases, crystal growth greatly benefited from micro-seeding. To limit vibration, crystallization trays were placed on foam padding.

### Data Collection and Structure Determination

X-ray diffraction data was collected at 100 K. Cocrystals were soaked in cryogenic solutions made by supplementing the exact precipitant solutions with 25 % glycerol except for the M^pro^ substrate structure C145A-NSP 7/8 where 20 % ethylene glycol was used. Crystallographic data was collected locally at the University of Massachusetts Chan Medical School X-Ray Crystallography Core facility and at the Brookhaven National Laboratory NSLS-II Beamline 17-ID-2 (FMX). In-house data collection was performed with a Rigaku MicroMax-007HF x-ray generator with either a Saturn944 or HyPix-6000HE detector. Diffraction data was indexed, integrated, and scaled using HKL2000 (HKL Research Inc.) or CrysAlis^pro^PX (Rigaku Corporation). NSLS-II collected diffraction intensities were automatically indexed, integrated, and scaled using XDS (39). All structures were solved using molecular replacement with PHASER (40). Model building and refinement were performed using Coot(41) and Phenix (42). The reference model used was PDB 7L0D (37). Prior to molecular replacement, the model was modified by removing all water, buffer, and cryogenic molecules as well as the small molecule inhibitor in the active site. To minimize reference model bias, 5 % of the data was reserved to calculate R_free_ (43). X-ray data collection parameters and refinement statistics are presented in Tables 1 and 2 in the supplementary data section.

### Structural analysis: hydrogen bonds, van der Waals calculations and the substrate envelope

The co-crystal structures contained either a M^Pro^ monomer or dimer in the asymmetric unit. For complexes with a dimer in the asymmetric unit, the protease chain with better electron density around the active site and substrate was chosen for analysis. The chain D was chosen for nsp4-nsp5, nsp5-nsp6, nsp8-nsp9, nsp9-nsp10; and chain C for nsp10-nsp11, nsp15-nsp16. Hydrogen bonds were determined using the show_contacts PyMOL Plugin with default parameters where the bond angle is between 63 and 180 degrees and the distance less than 4.0 A for any and 3.6 A for an ideal hydrogen bond between the proton and heavy atom.

Prior to van der Waals calculations, the crystal structures were prepared using the Schrodinger Protein Preparation Wizard (44). Hydrogen atoms were added, protonation states determined, and the hydrogen bonding network was optimized. A restrained minimization was performed using the OPLS2005 force field (45) within an RMSD of 0.3 Å. All crystallographic waters were retained during structure minimization. Interaction energies between the inhibitor and protease were estimated using a simplified Lennard-Jones potential, as previously described (46).

To generate the substrate envelope and other analyses, the cocrystal structures were superimposed using the carbon alpha atoms of active site residues 41, 144, 145, 163, 164 within one monomer. After superimposition, a Gaussian object map was generated for each substrate where the van der Waals volume was mapped onto a grid with a spacing of 0.5 A. The intersecting volumes of 4 substrates for all 126 possible combinations of the 9 substrates were calculated. Summation of these maps generated the consensus volume occupied by at least 4 of the substrates, which was used to construct the substrate envelope in PyMOL. A similar method was used to generate the product envelope, where the consensus of at least 4 products out of the 5 product cocrystal structures were determined.

As a second method to generate the substrate envelope, a custom python script was written to place a 3D grid with a spacing of 0.2 A at the active site and occupancy of each grid cell was counted in the 9 cocrystal structures. The grid cell was occupied when the van der Waals volume of a substrate atom was within the cell. The grid cells were given scores between 0 and 1 by normalizing the occupancy by the number of structures, and the resulting substrate envelope was visualized by coloring according to the calculated scores.

The figures were generated using Matplotlib (47), PyMOL and Maestro by Schrödinger LLC.

## Acknowledgements

This work was sponsored by Novartis Institutes for BioMedical Research. The author’s would like to acknowledge Elenore A. Wiggin, Winnie W. Mkandarwire and Vincent N. Azzolino for technical assistance.

