## Supplemental Materials for "Defining the Substrate Envelope of SARS-CoV-2 Main Protease to Predict and Avoid Drug Resistance"

#joint last authors

**Table S1:** Crystallization and refinement statistics of SARS-CoV2-M$^{pro}$-C145A in complex with substrate cleavage sites between nonstructural proteins (nsp).

| Substrate Complexes | **nsp4 - nsp5** | **nsp5 - NSP 6** | **nsp7- nsp8** | **nsp8- nsp9** | **nsp9- nsp10** | **nsp10- nsp11** |
|---|---|---|---|---|---|---|
| PDB ID | 7T70 | 7T8M | 7T8R | 7T9Y | 7TA4 | 7TA7 |
| **Data Collection** | | | | | | |
| Location | Home source | Home source | Home source | Home source | NSLS-II | Home source |
| Resolution range (Å) | 29.82 - 2.354 (2.438 - 2.354) | 23.92 - 1.6 (1.657 - 1.6) | 24.37 - 1.74 (1.802 - 1.74) | 28.93 - 2.18 (2.258 - 2.18) | 27.20 - 1.781 (1.845 - 1.781) | 23.33 - 2.281 (2.362 - 2.281) |
| Space group | $P2_12_12_1$ | $P2_12_12_1$ | C2 | $P2_12_12_1$ | C2 | $P12_11$ |
| a, b, c (Å) | 67.4, 99.7, 101.8 | 67.6, 99.7, 103.4 | 98.6, 80.0, 51.9 | 67.2, 98.9, 101.6 | 101.8, 78.8, 103.8 | 46.7, 96.4, 67.4 |
| $\alpha$, β, γ (º) | 90, 90, 90 | 90, 90, 90 | 90, 114.5, 90 | 90, 90, 90 | 90, 115.5, 90 | 90, 103.2, 90 |
| Total reflections | 202351 (17367) | 466655 (22545) | 270230 (16820) | 219740 (17048) | 135086 (12711) | 95173 (8250) |
| Unique reflections | 28782 (2756) | 85584 (7267) | 37551 (3732) | 35387 (3381) | 69743 (6769) | 26383 (2538) |
| Multiplicity | 7.0 (6.3) | 5.5 (3.1) | 7.2 (4.5) | 6.2 (5.0) | 1.9 (1.9) | 3.6 (3.2) |
| Completeness (%) | 98.74 (96.23) | 92.38 (79.46) | 98.53 (99.28) | 98.24 (95.64) | 98.48 (96.40) | 99.41 (96.14) |
| Average I/$\sigma$ | 12.96 (3.17) | 22.08 (2.51) | 22.39 (2.16) | 14.33 (3.75) | 9.46 (1.71) | 9.62 (3.04) |
| Wilson B-factor | 26.08 | 10.18 | 21.13 | 25.82 | 21 | 29.14 |
| $R_{merge}{}^a$ | 0.1284 (0.5863) | 0.05001 (0.4511) | 0.06852 (0.5145) | 0.08742 (0.4391) | 0.05304 (0.4258) | 0.09104 (0.3705) |
| $CC_{1/2}$ | 0.997 (0.882) | 0.999 (0.813) | 0.999 (0.925) | 0.998 (0.897) | 0.998 (0.709) | 0.995 (0.85) |
| **Refinement** | | | | | | |
| $R_{factor}{}^c$ | 0.1715 (0.2172) | 0.1605 (0.2020) | 0.1845 (0.2554) | 0.1794 (0.2334) | 0.1750 (0.2380) | 0.2048 (0.2505) |
| $R_{free}{}^d$ | 0.2199 (0.2930) | 0.1971 (0.2396) | 0.2278 (0.2849) | 0.2363 (0.3140) | 0.2163 (0.2603) | 0.2545 (0.3131) |
| RMSD[b] in: | | | | | | |
| Bond lengths (Å) | 0.002 | 0.016 | 0.018 | 0.003 | 0.008 | 0.002 |
| Bond angles (º) | 0.50 | 1.47 | 1.63 | 0.59 | 0.97 | 0.40 |
| Ramachandran: | | | | | | |
| Favored (%) | 96.49 | 97.75 | 97.08 | 97.41 | 98.71 | 96.15 |
| Allowed (%) | 3.51 | 2.09 | 2.92 | 2.59 | 1.29 | 3.85 |
| Outliers (%) | 0 | 0.16 | 0 | 0 | 0 | 0 |
| Rotamer outliers (%) | 1.49 | 0.19 | 1.12 | 0.58 | 0.57 | 0.6 |
| B-factors: | | | | | | |
| Average | 29.05 | 15.91 | 28.23 | 28.65 | 26.42 | 32.8 |
| Macromolecules | 27.82 | 14.07 | 26.95 | 28.11 | 24.96 | 32.6 |
| Solvent | 36.23 | 26.65 | 36.85 | 34.25 | 35.81 | 36.33 |

[a]$R_{sym} = \Sigma \mid I - \langle I \rangle \mid / \Sigma I$, where $I$ = observed intensity, $\langle I \rangle$ = average intensity over symmetry equivalent.
[b]RMSD, root mean square deviation.
[c]$R_{factor} = \Sigma \mid\mid F_o \mid - \mid F_c \mid\mid / \Sigma \mid F_o \mid$.
[d]$R_{free}$ was calculated from 5% of reflections, chosen randomly, which were omitted from the refinement process.
Statistics for the highest-resolution shell are shown in parentheses.

**Table 1 continued.**

| Substrate Complexes | **nsp12- nsp13** | **nsp13 - nsp14** | **nsp15- nsp16** |
|---|---|---|---|
| PDB ID | 7TB2 | 7TBT | 7TC4 |
| **Data Collection** | | | |
| Location | Home source | Home source | Home source |
| Resolution range (Å) | 24.37 - 1.8 (1.864 - 1.8) | 22.86 - 2.452 (2.539 - 2.452) | 25.77 - 1.94 (2.009 - 1.94) |
| Space group | C2 | C2 | $P12_11$ |
| a, b, c (Å) | 99.0, 79.4, 51.9 | 98.0, 78.0, 51.8 | 54.8, 102.8, 67.6 |
| $\alpha$, β, γ (º) | 90, 114.66, 90 | 90, 114.40, 90 | 90, 92.59, 90 |
| Total reflections | 67722 (6722) | 36712 (3394) | 262729 (17543) |
| Unique reflections | 33884 (3375) | 12692 (1226) | 55231 (5476) |
| Multiplicity | 2.0 (2.0) | 2.9 (2.8) | 4.8 (3.2) |
| Completeness (%) | 98.90 (99.14) | 95.40 (94.94) | 99.83 (99.82) |
| Average $I/\sigma$ | 19.45 (3.36) | 9.58 (3.89) | 12.74 (1.53) |
| Wilson B-factor | 20.08 | 34.97 | 18.41 |
| $R_{merge}$[a] | 0.02335 (0.1794) | 0.06682 (0.2865) | 0.09075 (0.5802) |
| $CC_{1/2}$ | 0.999 (0.942) | 0.995 (0.939) | 0.995 (0.671) |
| **Refinement** | | | |
| $R_{factor}$[c] | 0.1876 (0.2495) | 0.2102 (0.2831) | 0.1821 (0.2427) |
| $R_{free}$[d] | 0.2179 (0.2973) | 0.2664 (0.3582) | 0.2267 (0.2941) |
| RMSD[b] in: | | | |
| Bond lengths (Å) | 0.011 | 0.002 | 0.01 |
| Bond angles (º) | 1.11 | 0.52 | 1.03 |
| Ramachandran: | | | |
| Favored (%) | 98.38 | 97.11 | 97.58 |
| Allowed (%) | 1.62 | 2.89 | 2.26 |
| Outliers (%) | 0 | 0 | 0.16 |
| Rotamer outliers (%) | 0 | 3.08 | 0.75 |
| B-factors: | | | |
| Average | 23.96 | 38.95 | 23.29 |
| Macromolecules | 22.73 | 38.89 | 22.41 |
| Solvent | 33.07 | 40.02 | 30.59 |

[a]$R_{sym} = \Sigma \mid I - \langle I \rangle \mid / \Sigma I$, where $I$ = observed intensity, $\langle I \rangle$ = average intensity over symmetry equivalent.
[b]RMSD, root mean square deviation.
[c]$R_{factor} = \Sigma \| F_o| - |F_c\| / \Sigma |F_o|$.
[d]$R_{free}$ was calculated from 5% of reflections, chosen randomly, which were omitted from the refinement process.
Statistics for the highest-resolution shell are shown in parentheses.

**Table 2:** Crystallization and refinement statistics of SARS-CoV2-M$^{pro}$-C145A in complex with proteolyzed cleavage sites between nonstructural proteins (nsp).

| Product Complexes | **nsp 4** | **nsp 5** | **nsp 6** | **nsp 7** | **nsp 8** | **nsp 10** |
|---|---|---|---|---|---|---|
| PDB ID | 7MB4 | 7MB5 | 7MB6 | 7MB7 | 7MB8 | 7MB9 |
| **Data Collection** | | | | | | |
| Location | NSLS-II | NSLS-II | NSLS-II | NSLS-II | NSLS-II | NSLS-II |
| Resolution range (Å) | 29.6 - 1.83 (1.895 - 1.83) | 28.76 - 1.6 (1.657 - 1.6) | 28.31 - 2.211 (2.29 - 2.211) | 29.09 - 2.02 (2.092 - 2.02) | 29.49 - 1.624 (1.682 - 1.624) | 29.24 - 1.814 (1.879 - 1.814) |
| Space group | $P12_11$ | $P2_12_12_1$ | C2 | C2 | $P12_11$ | $P12_11$ |
| a, b, c (Å) | 69.7, 100.6, 99.3 | 67.9, 100.1, 103.1 | 124.9, 80.8, 62.7 | 99.4, 79.4, 52.1 | 67.8, 103.0, 104.3 | 54.2, 97.9, 67.5 |
| $\alpha$, β, γ (º) | 90, 104.2, 90 | 90, 90, 90 | 90, 91.3, 90 | 90, 114.5, 90 | 90, 101.4, 90 | 90, 102.5, 90 |
| Total reflections | 230216 (22187) | 180287 (17298) | 61772 (5885) | 47317 (4338) | 344159 (28355) | 120841 (10414) |
| Unique reflections | 115488 (11277) | 90251 (8704) | 31114 (3000) | 23874 (2236) | 174513 (15578) | 61128 (5401) |
| Multiplicity | 2.0 (2.0) | 2.0 (2.0) | 2.0 (2.0) | 2.0 (1.9) | 2.0 (1.8) | 2.0 (1.9) |
| Completeness (%) | 98.70 (97.32) | 96.72 (94.07) | 99.35 (95.45) | 98.30 (93.04) | 98.66 (88.38) | 98.21 (86.91) |
| Average I/$\sigma$ | 12.51 (2.46) | 12.65 (2.34) | 12.72 (1.89) | 10.18 (1.90) | 11.69 (1.75) | 10.12 (1.44) |
| Wilson B-factor | 24.55 | 18.18 | 44.29 | 39.68 | 17.49 | 30.37 |
| $R_{merge}$[a] | 0.02846 (0.2512) | 0.02716 (0.2411) | 0.03287 (0.3806) | 0.03295 (0.2949) | 0.0347 (0.375) | 0.03511 (0.3266) |
| $CC_{1/2}$ | 0.999 (0.871) | 0.999 (0.844) | 0.999 (0.719) | 0.998 (0.882) | 0.999 (0.813) | 0.998 (0.839) |
| **Refinement** | | | | | | |
| $R_{factor}$[c] | 0.1770 (0.2443) | 0.1643 (0.2333) | 0.2226 (0.3122) | 0.1900 (0.3084) | 0.1686 (0.2506) | 0.1989 (0.3138) |
| $R_{free}$[d] | 0.2198 (0.2730) | 0.1838 (0.2500) | 0.2720 (0.3951) | 0.2261 (0.3490) | 0.1951 (0.2853) | 0.2276 (0.3457) |
| RMSD[b] in: | | | | | | |
| Bond lengths (Å) | 0.019 | 0.009 | 0.003 | 0.003 | 0.013 | 0.013 |
| Bond angles (º) | 1.4 | 1.18 | 0.62 | 0.64 | 1.08 | 1.03 |
| Ramachandran: | | | | | | |
| Favored (%) | 97.31 | 98.21 | 95.91 | 97.37 | 98.45 | 96.56 |
| Allowed (%) | 2.69 | 1.63 | 4.09 | 2.63 | 1.47 | 3.27 |
| Outliers (%) | 0 | 0.16 | 0 | 0 | 0.08 | 0.16 |
| Rotamer outliers (%) | 0 | 0.39 | 0.21 | 0.39 | 0.19 | 0 |
| B-factors: | | | | | | |
| Average | 30.10 | 22.80 | 53.41 | 50.09 | 23.91 | 36.47 |
| Macromolecules | 29.22 | 21.26 | 53.5 | 50.01 | 21.94 | 36 |
| Solvent | 37.73 | 33.87 | 49.8 | 51.49 | 35.52 | 41.74 |

[a]$R_{sym} = \Sigma \mid I - \langle I \rangle \mid / \Sigma I$, where $I$ = observed intensity, $\langle I \rangle$ = average intensity over symmetry equivalent.
[b]RMSD, root mean square deviation.
[c]$R_{factor} = \Sigma \| F_o| - |F_c\| / \Sigma |F_o|$.
[d]$R_{free}$ was calculated from 5% of reflections, chosen randomly, which were omitted from the refinement process.
Statistics for the highest-resolution shell are shown in parentheses

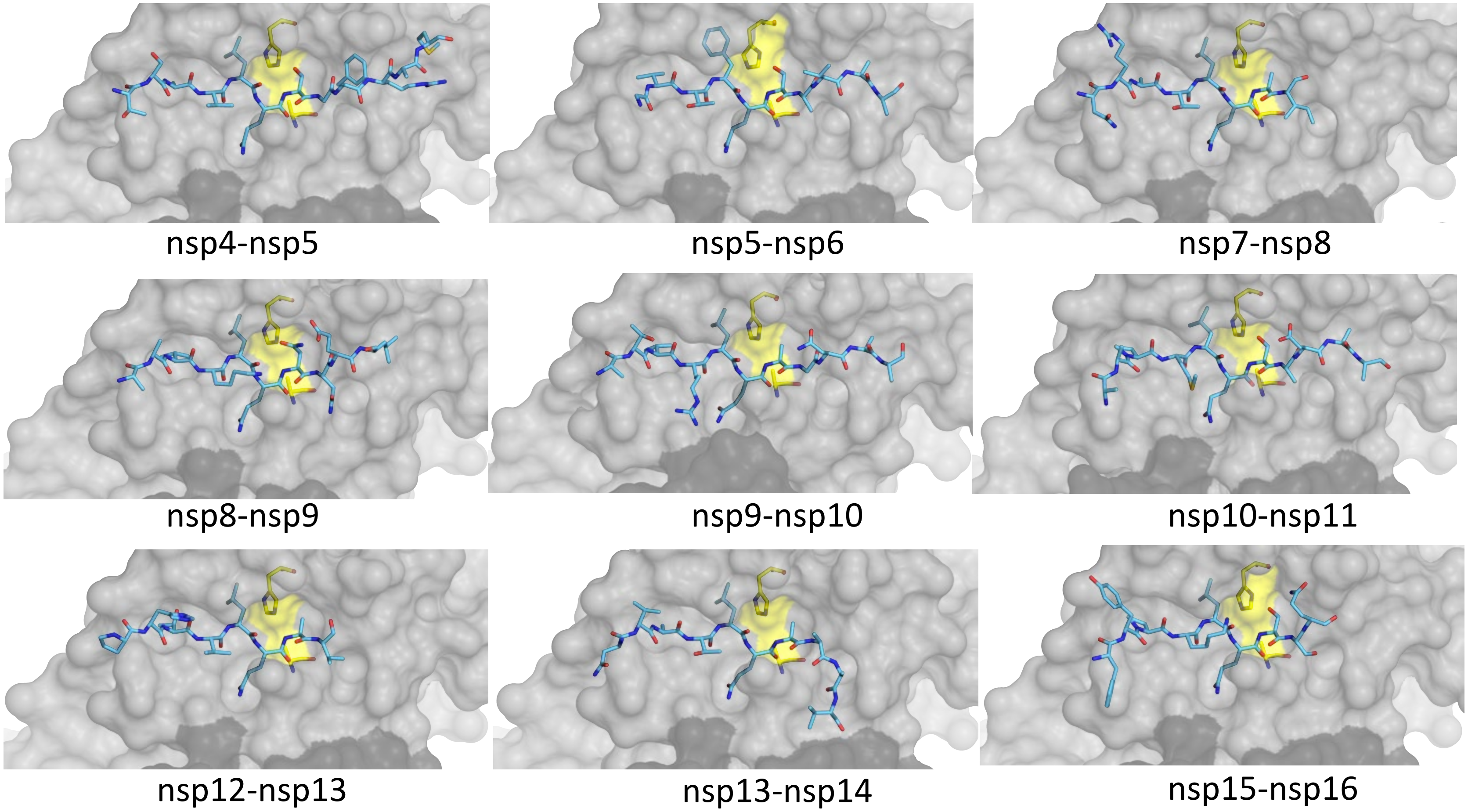


**Figure S1. Substrates bound at the active site of $M^{pro}$ in the cocrystal structures determined.** The peptide is depicted as cyan sticks and the catalytic dyad is colored yellow. The panels show close-up view of the active sites, with the protease in gray surface representation, similar to Figure 1C of the main manuscript.

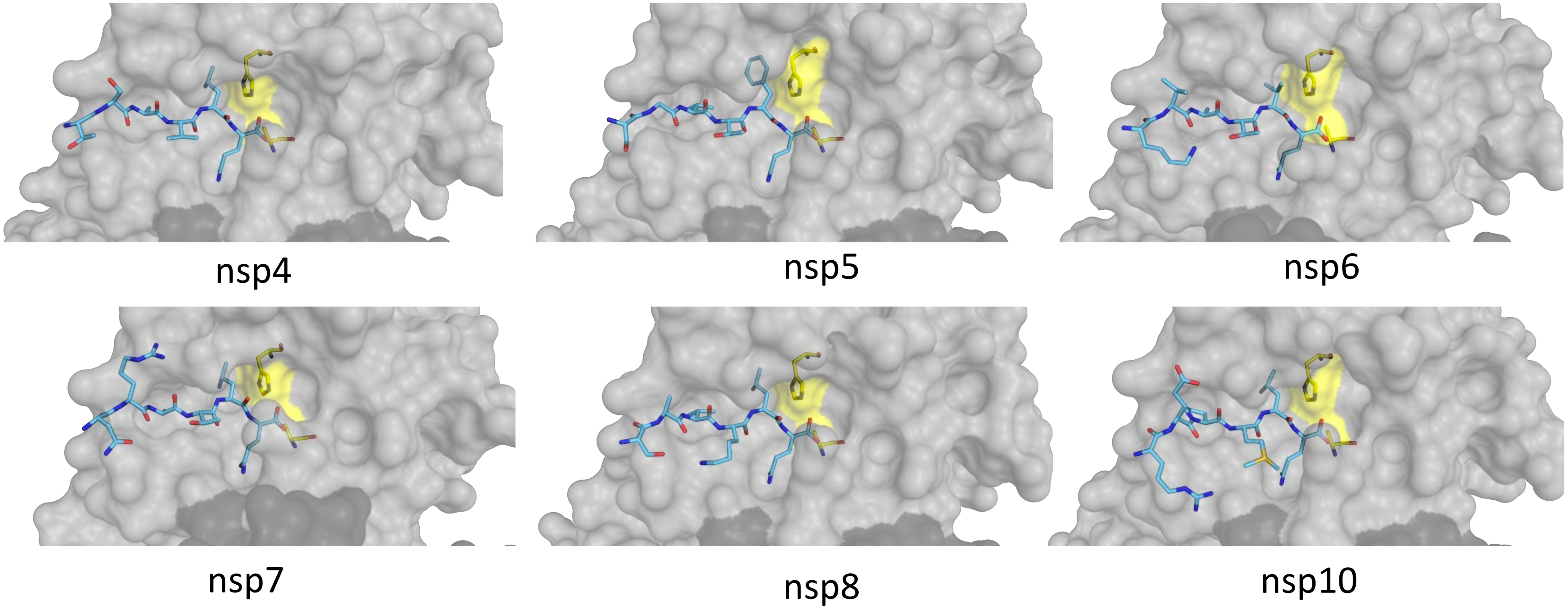


**Figure S2. Substrates cleaved with the N-terminal product bound at the active site of $M^{pro}$ in the cocrystal structures determined.** The peptide is depicted as cyan sticks and the catalytic dyad is colored yellow. The panels show close-up view of the active sites, with the protease in gray surface representation.

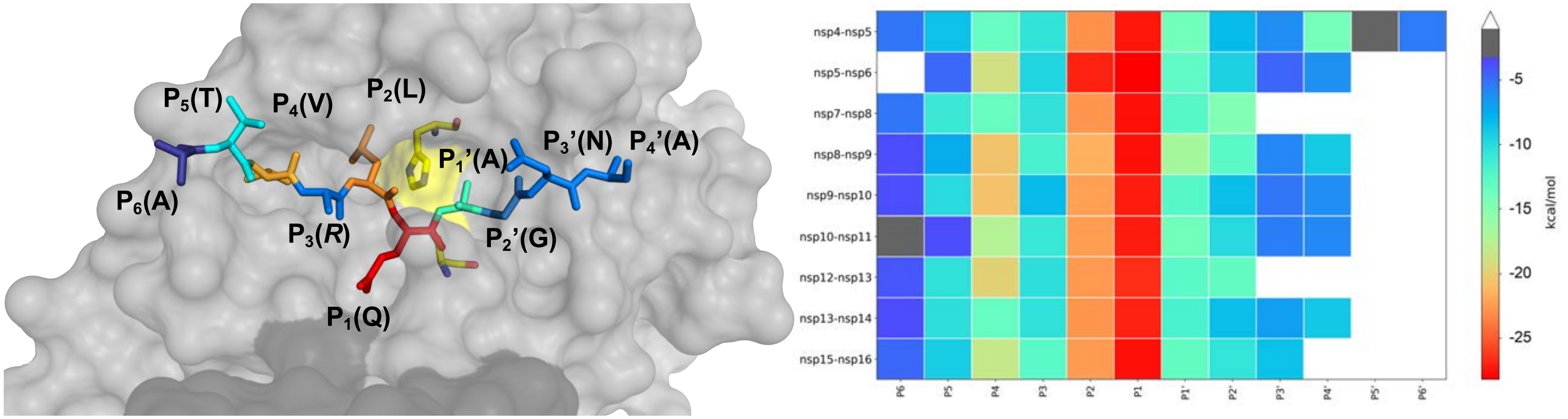


**Figure S3. Substrate vdW interactions with the active site of M$^{pro}$ in the cocrystal structures.** Similar to Figure 3A of the main manuscript, the peptide is depicted as sticks and the catalytic dyad is colored yellow. The substrate residues are colored from blue to red according to increasing van der Waals interactions with the protease. The heat map shows per residue interactions for all 9 substrate cocrystal structures.

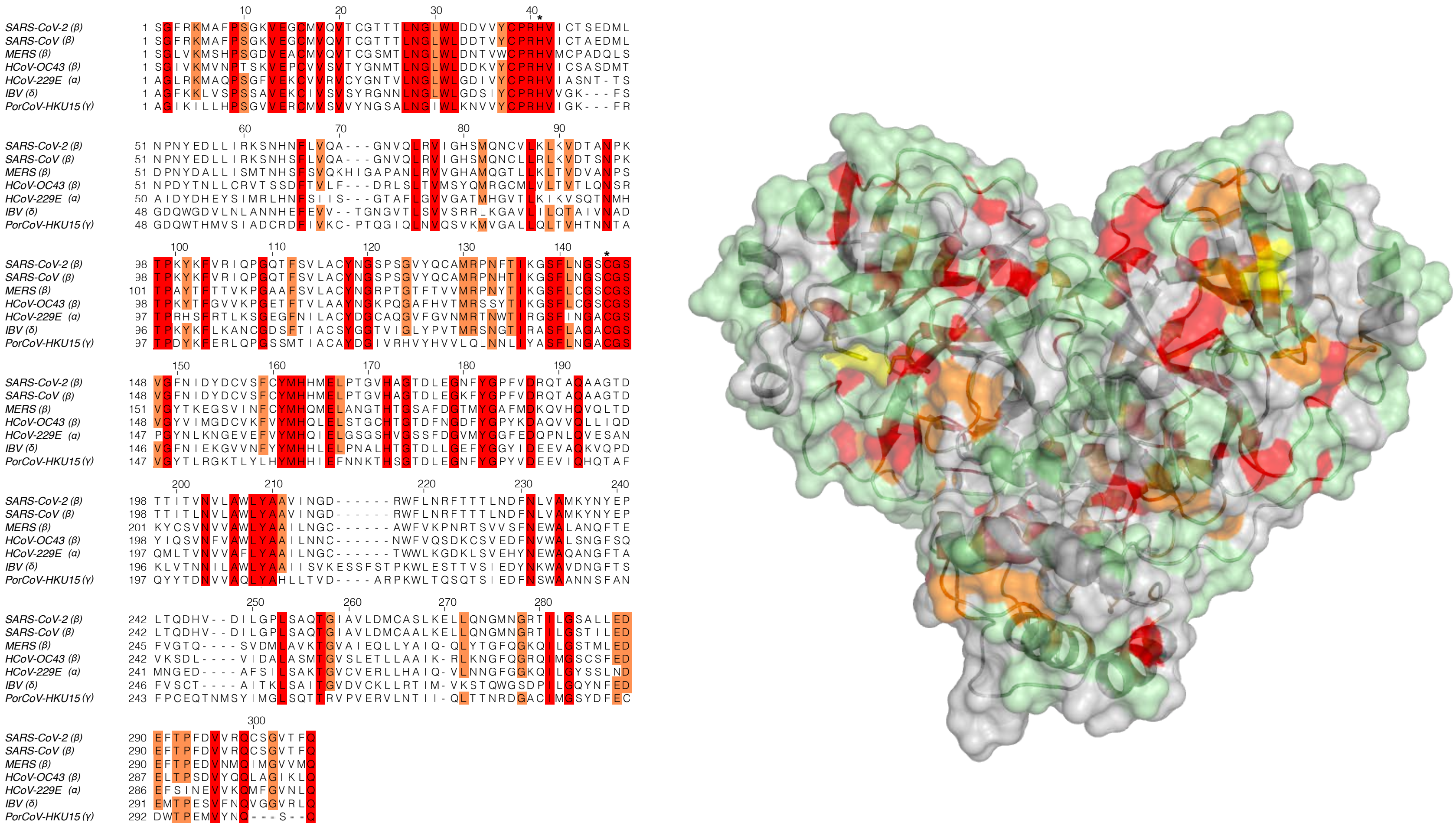


**Figure S4. Sequence conservation of M$^{pro}$.** Alignment of 7 diverse sequences coronaviral sequences of M$^{pro}$ -- Beta: SARS-CoV-2, SARS-CoV, MERS, Human coronavirus OC43 (HCoV-OC43); alpha: Human coronavirus 229E (HCoV-229E); delta: Avian coronavirus IBV; gamma: Porcine Coronavirus HKU15(PorCoV-HKU15) (left) and the amino acid sequence conservation of M$^{pro}$ betweenthe 7 coronaviral species depicted on the structure where surface residues identical are shown in all 7 (red), 5-6 of 7 (orange), 3-4 of 7 (green) and less than 3 (highly variable; gray) sequences are indicated by color (right).

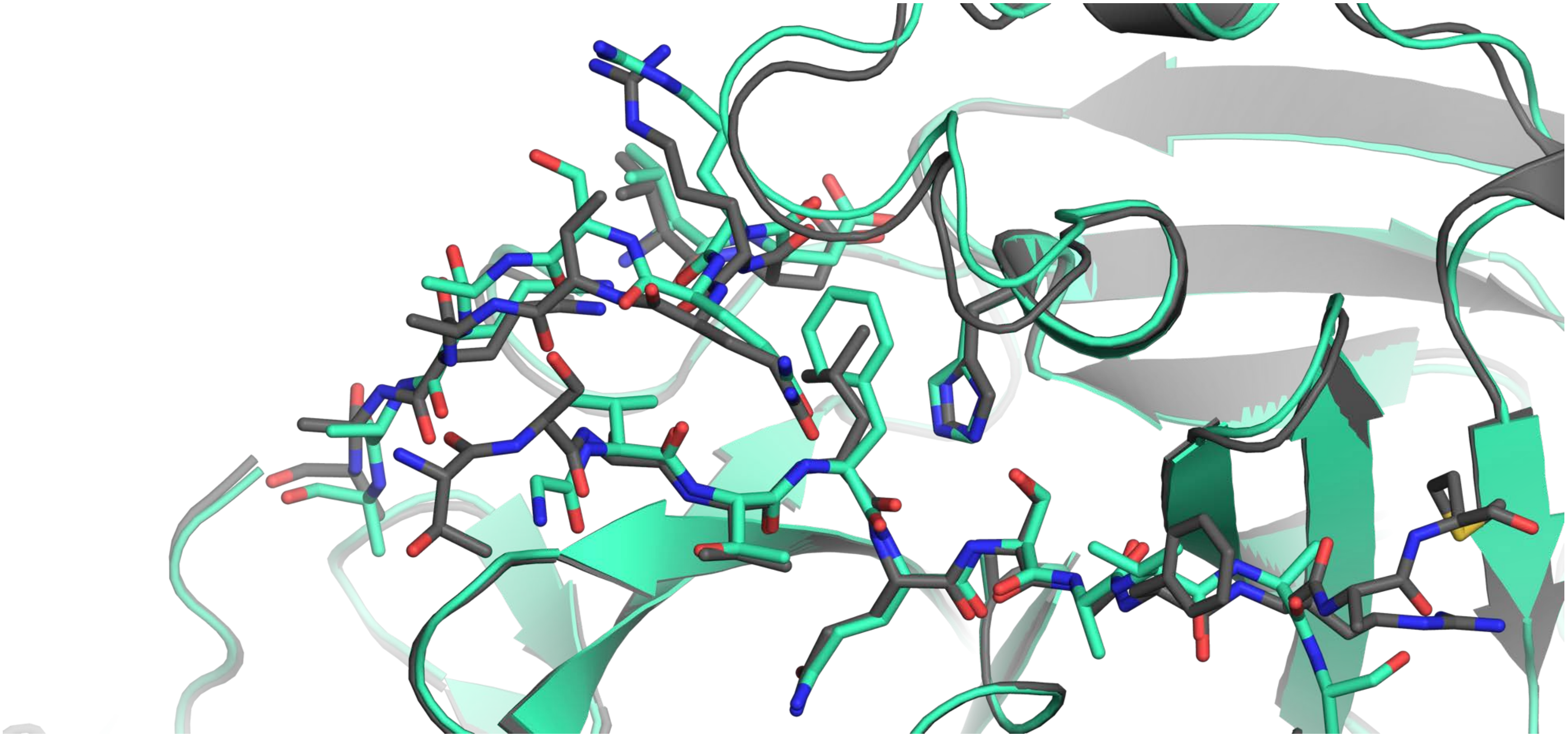

**Figure S5. Comparison of the variable 187-192 loop in substrate–M^pro cocrystal structures.** The substrate and the loop residues depicted as sticks as well as the catalytic His side chain.  Nsp5-nsp6 with a Phe at P2 is the most divergent (in green), shown in comparison with nsp4-nsp5

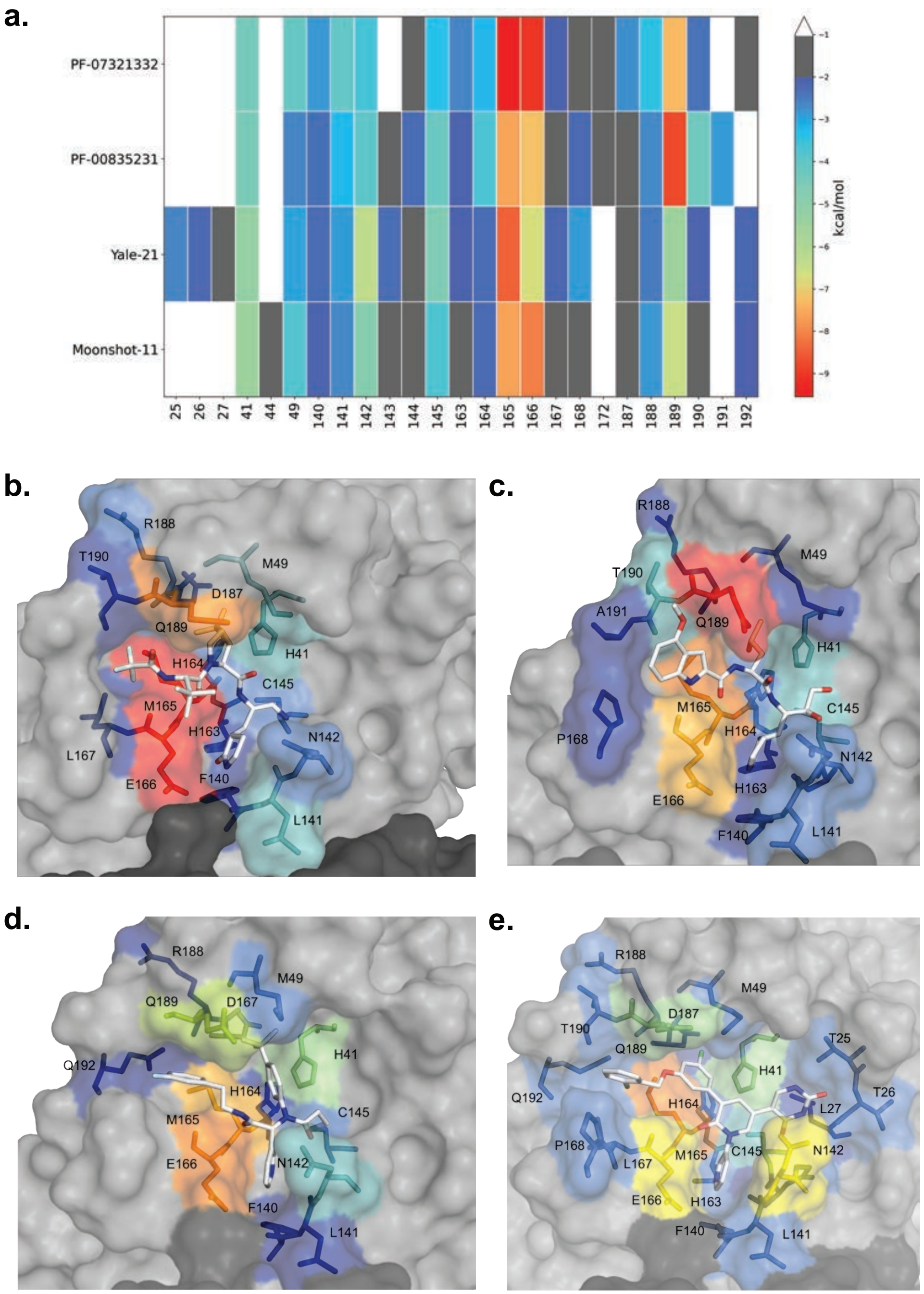
a.
PF-07321332
PF-00835231
Yale-21
Moonshot-11
25
26
27
41
44
49
140
141
142
143
144
145
163
164
165
166
167
168
172
187
188
189
190
191
192
kcal/mol
b.
R188
T190
M49
D187
Q189
H164
H41
C145
M165
H163
N142
L167
E166
F140
L141
c.
R188
M49
T190
A191
Q189
H41
M165
C145
P168
H164
N142
H163
E166
F140
L141
d.
R188
M49
Q189
D167
H41
Q192
H164
C145
M165
E166
N142
F140
L141
e.
R188
M49
D187
T190
T25
Q189
H41
T26
Q192
H164
L27
P168
L167
M165
C145
N142
E166
H163
F140
L141

**Figure S6. Inhibitor Van der Waals** heatmap interactions on the active site $M^{pro}$ **(A)** colored by extent of van der Waals **(B)** PF-07321332 (25), (**C)** PF-00835231 (26), PDBID: 6XHM **(D)** Noncovalent potent compound 21 (27), PDBID: 7L13 **(E)** Moonshot compound 11 (28), PDBID: 7NW2.
